# Intraspecific variation promotes species coexistence and trait clustering through higher order interactions

**DOI:** 10.1101/494757

**Authors:** Gaurav Baruah, Robert John

**Affiliations:** University of Zurich, Department of Evolutionary biology and Environmental studies; Indian Institute of Science Education and Research, Kolkata

**Author notes:** **Data Accessibility Statement:** Data and code will be made available in Dryad repository.

**Keywords:** higher-order interactions, intraspecific variation, eco-evolutionary dynamics, species coexistence, trait clustering, stability

## Abstract

Ecological and evolutionary effects of individual variation on species coexistence remains unclear. Competition models for coexistence have emphasized species-level differences in pairwise interactions, and invoked no role for intraspecific variation. These models show that stronger competitive interactions result in smaller numbers of coexisting species. However, the presence of higher-order interactions (HOIs) among species appears to have a stabilizing influence on communities. How species coexistence is affected in a community where both pairwise and higher-order interactions are pervasive is not known. Furthermore, the effect of individual variation on species coexistence in complex communities with pairwise and HOIs remains untested. Using a Lotka-Volterra model, we explore the effects of intraspecific variation on the patterns of species coexistence in a competitive community dictated by pairwise and HOIs. We found that HOIs greatly stabilize species coexistence across different levels of strength in competition. Notably, high intraspecific variation promoted species coexistence, particularly when competitive interactions were strong. However, species coexistence promoted by higher levels of variation was less robust to environmental perturbation. Additionally, species’ traits tend to cluster together when individual variation in the community increased. We argue that individual variation can promote species coexistence by reducing trait divergence and attenuating the inhibitory effects of dominant species through HOIs

## Introduction

Explanations for multi-species coexistence in ecological communities have largely been sought at the species level by emphasizing average life history differences among species driven by competitive interactions or trade-offs (Clark 2010*a*; Gravel et al. 2011; Violle et al. 2012; Kraft et al. 2015; Valladares et al. 2015; Letten et al. 2017; Wittmann and Fukami 2018). Such differences among species in multiple ecological dimensions could minimize niche overlap and promote long-term species coexistence (Clark et al. 2010; Barabas and D’Andrea 2016; Barabás et al. 2016). However, clear niche differences among species has rarely been found, and in fact large numbers of species appear to compete for only a small number of limiting resources, giving rise to a paradox (Hutchinson 1961; Laird and Schamp 2006; Shoresh et al. 2008; Li and Chesson 2016; Letten et al. 2018). Many species coexist despite little measurable difference in demographic or resource-based niches (Condit et al. 2006). So, although there are strong theoretical arguments that average differences among species can account for species coexistence, empirical support has remained scanty.

Classical competition models of coexistence consider interactions among species pairs that require precise parameter trade-offs to stabilize communities or to limit the strength of competition in accordance with the competitive exclusion principle (Barabás and Meszéna 2009; Barabás et al. 2016). The implausibility of highly structured competitive relationships in species-rich communities has prompted models of coexistence based on ecological equivalence rather than life historical differences (Hubbell 2006; Rosindell et al. 2011; Segura et al. 2011). Theoretical studies with competition models further show that any stability achieved through structured pairwise competitive interactions can be disrupted by random interactions among species (Allesina and Levine 2011; Bairey et al. 2016; Barabás et al. 2016). The number of coexisting species then declines inversely with the strength of interactions among species pairs (Bairey et al. 2016). Interaction strength therefore places an upper bound on the numbers of coexisting species, implying that strong pairwise competitive interactions alone cannot promote species coexistence in a large community.

Interactions among species are not always constrained to species pairs but can involve higher-order combinations (Wilson 1992; Laird and Schamp 2006; Bairey et al. 2016; Grilli et al. 2017; Mayfield and Stouffer 2017; Terry et al. 2017), where interactions between a species pair is modulated by a third or more species (Fig. 1). In an ecological system where communities are structured by pairwise interactions indirect or higher-order effects may alter these interactions and restructure communities (Terhorst et al. n.d.; Levine et al. 2017). For example, a species that is a superior competitor for a given resource can inhibit an inferior competitor for the same resource, but a third species may modulate the strength of this inhibition without affecting either of the two competitors directly (Bairey et al. 2016). Such attenuation of the pairwise inhibitory effect can be density-mediated or trait-mediated, and can lead to qualitatively different community dynamics compared to pure pairwise interactions. The importance of such higher-order interactions has been recognised (Levine et al. 2017), but the singular focus of coexistence studies on average species level differences has meant that few investigations have been undertaken.

**Figure 1.**
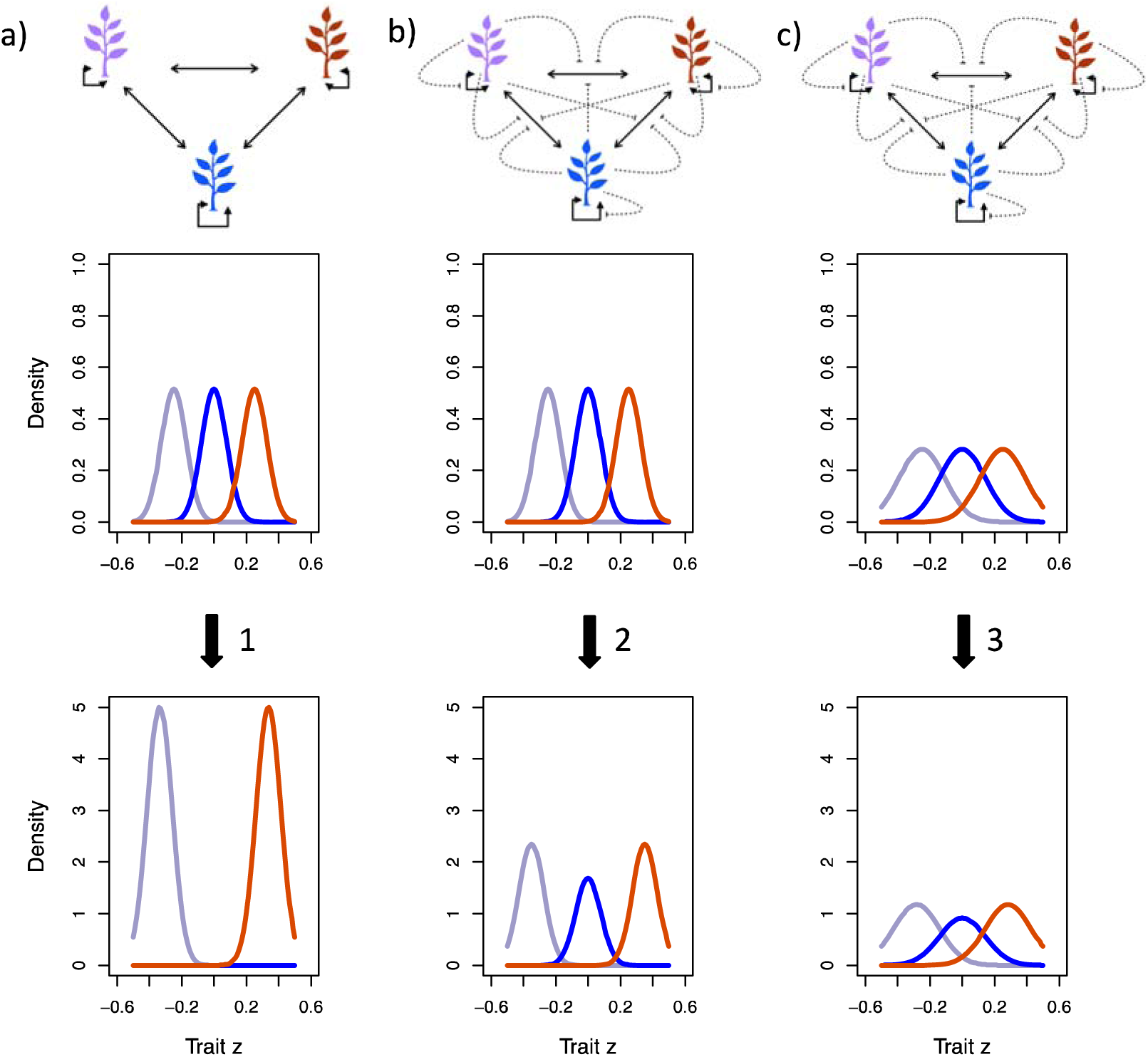
An example image of how intraspecific trait variation in communities with (a) pairwise competitive interactions (black arrows) and (b), (c) higher-order interactions (dashed-arrows) can affect trait patterning and coexistence. (a) Three different species (Red, blue and orange) are spaced along a trait axis with high variation. Interactions between the three species are inherently pairwise. Starting with purely pairwise competitive communities’ initial high intraspecific variation will ultimately eventually lead (1) to competitive exclusion of the red species. In addition, the other two remaining species (blue and orange) will space themselves far apart minimizing trait overlap and leading to the emergence of what is called the ‘limiting similarity’ principle. However, with the introduction of higher-order interactions, low levels of intraspecific variation (b) will also lead to (2) species minimizing trait overlap but leading to all species coexisting. However, (c) high intraspecific variation and with higher-order interactions will lead to (3) more trait overlap as well as coexistence of all the three species.

A further consequence of the emphasis on species-level differences is that within-species or individual level variation has largely been ignored (Siefert 2012; Hart et al. 2016). Observations that variation within species often exceeds the differences in species-level averages have inspired much theoretical and empirical research (Barabas and D’Andrea 2016; Barabás et al. 2016; Hart et al. 2016; Hausch et al. 2018). Intraspecific variation can have both ecological and evolutionary effects on competitive interactions, and therefore on species coexistence. For example, intraspecific trait variation can hamper species coexistence by increasing competitive ability, niche overlap and even-spacing among species (Barabas and D’Andrea 2016), or by altering competitive outcomes through non-linear averaging of performances (Hart et al. 2016). There is equally compelling evidence that intraspecific variation promotes species coexistence, mainly through disruption of interspecific competitive abilities and obscuring the effect of strongly competitive individuals in a community (Bolnick et al. 2011). Experimental work has shown that while intraspecific variation allows a community to be resilient to invaders, may create the opportunity for competitive exclusion among strong competitors (Hausch et al. 2018). Empirical studies have consistently found that most of the variation in plant life histories lies within individuals rather than species. Tree growth rates vary remarkably within individuals (Rüger et al. 2011, Clark 2010), and variation may be driven by several factors including light, soil resources, herbivores, and pathogens, that affect growth. High intraspecific variation in leaf economics such as specific leaf area, leaf N and P, may be driven by how individuals tap the variation along these dimensions (Meziane and Shipley 1999; Vasseur et al. 2012). In multi-species communities diversity appears to be positively correlated with intraspecific variability (Fridley and Grime 2010). Conversely, the loss of individual trait variation in plant communities could lead to increases in susceptibility to plant invasions (Crutsinger et al. 2007). Given that high levels of intraspecific trait variation within communities appears to be more a rule than an exception, the combined influence of intraspecific variation, and pairwise and higher-order species interactions on diversity and community structure merits detailed investigation.

Theoretical research on species coexistence has largely focused on the importance of higher-order species interactions and intraspecific variation separately. The effect of intraspecific trait variation and eco-evolutionary dynamics on structuring large communities where both pairwise and higher-order interactions dominate a community is unknown. Purely pairwise interactions in a community may lead to even trait spacing when intraspecific variation is high. Consequently, competitive exclusion of inferior species in a large community becomes inevitable (Fig. 1). However, a community dominated by both pairwise and higher-order interactions could lead to less-even spacing of species in a trait axis and result in trait clustering. This could occur because with high intraspecific variation present in the community, higher-order interactions could significantly alleviate and stabilize the negative pairwise interactions that lead to distinct spacing in the first place. The link between HOIs and intraspecific variation therefore appears critical to understand coexistence in species-rich communities.

Here in this study, we examine the importance of higher order interactions and intraspecific variation in structuring species coexistence and trait patterning. We do this using a modified Lotka-Volterra modelling approach, where the dynamics of the whole community is mediated both by pairwise competitive interactions as well as higher-order three-way interactions. Specifically, we model a one-dimensional quantitative trait that contributes to the competitive ability of species interacting in the community. We show that in the presence of higher-order interactions, high intraspecific variation across different levels of strength in competition leads to significantly greater numbers of species coexisting in a community than when individual variation is low. We show analytically and with model simulations that intraspecific variation not only contributes to species coexistence, but also stabilizes the community to external perturbation. In addition, our analyses reveal that intraspecific variation in a community where higher-order interactions dictates dynamics leads to stable trait clustering. Our study links the recent ecological studies of higher-order interactions with eco-evolutionary dynamics and intraspecific variation.

## 2. Methods and Models

### 2.1 Community model with pairwise interactions

In our community model, we consider species competing with each other in a one-dimensional trait axis, where a species’ competitive ability is determined by a one-dimensional quantitative trait *z*. Individuals of a species vary along the competitive trait *z* of interest such that the distribution of the primary trait *z* is normally distributed with mean *u*_*i*_ for species *i* and variation given by 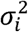. Under such conditions, the dynamics of a species *i* is given by Lotka-Volterra equations as (Barabas and D’Andrea 2016):

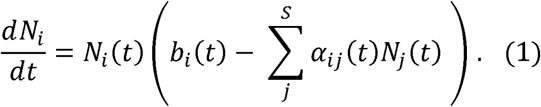

And the dynamics of the mean competitive trait *u*_*i*_ is given by:

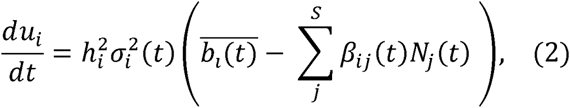

Where *α*_*ij*_(*t*) describes the pairwise competition coefficient of species *i* with species *j* at any time *t*. This competition coefficient derives directly from Gaussian competition kernel (See appendix 2). If the two species are similar to each other in terms of their average trait value *u*, then competition between them is stronger than when they are farther apart in the trait axis; 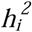 is the heritability of species *i, b* _*i*_(*t*) describes the growth rate of the species *i* in the absence of any competition which is determined by where they lie in the trait axis 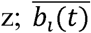 describes the growth of the trait and *β*_*ij*_ (*t*) quantifies the evolutionary pressure on the trait *z* of species *i* due to competition with the species *j* in the community (this has been derived in Barabas et al, 2016).

### 2.2 Community model with higher-order interactions

The above equations 1 and 2, captures the eco-evolutionary dynamics of a multispecies community where pairwise interactions dominate community dynamics. It is still plausible that such a community exhibits higher-order interactions than just between pairs of species. In extension to the above model, we include density-mediated three-way higher-order interactions where density of a third species influences pairwise competitive interactions. Under these circumstances, the equations become (see appendix 2):

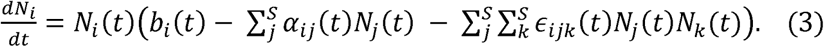

And the dynamics of the competitive trait *u*_*i*_ is given by:

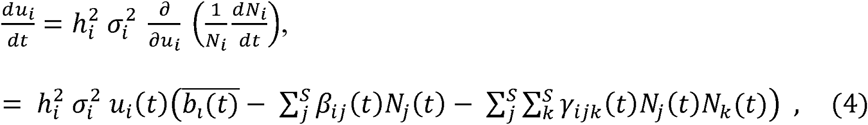

where *ϵ*_*ijk*_ (*t*) gives the 3-way interactions that are density mediated HOIs (*ϵ*_*ijk*_ is termed inter-specific HOIs and *ϵ*_*iii*_ *ϵ*_*iij*_ termed as intraspecific HOIs) (Mayfield and Stouffer 2017; Letten and Stouffer 2019); *γ*_*ijk*_ denotes 3-way interactions (*γ*_*ijk*_ as interspecific evolutionary effects; *γ*_*iii*_ *γ*_*iij*_ as intraspecific evolutionary HOI effects) affecting evolutionary dynamics of mean trait *u* for species *i*. Similar to the pairwise Gaussian interaction kernel, the three way interaction remains Gaussian with a third species *k* influencing the interaction between the two species *i* and *j* given as (see appendix 2):

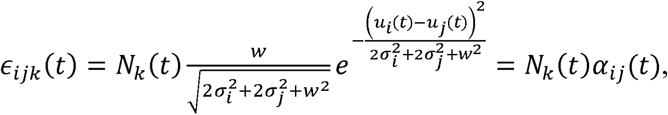

And, *γ*_*ijk*_ (*t*) can be written as (appendix 2):

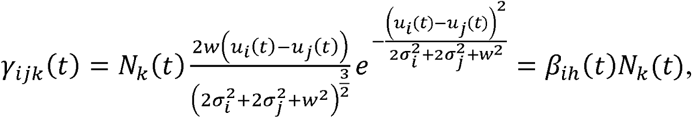

Where, *ϵ*_*ijk*_ (*t*)and *γ*_*ijk*_ (*t*) are three-dimensional tensors of size (S x S x S), where S is the total number of species in the community. Thus we can formally define intraspecific HOIs as *ϵ*_*iik*_ (*t*) and *ϵ*_*iii*_ (*t*) and interspecific HOIs as 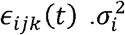 and 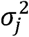 are the intraspecific trait variation for species *i* and species *j* respectively; *w*^2^ is the width of the competition kernel which is Gaussian (see appendix 2); *u*_*i*_(*t*) is the average trait value of species *i* and *u*_*j*_(*t*) is the average trait value for species *j*. Thereby, eco-evolutionary dynamics in this purely competitive community is dominated not only by pairwise trait-based competition but also by three-way higher-order interactions. In such a case, eco-evolutionary dynamics might deviate from dynamics dominated by purely pairwise competitive coefficients as in (Barabas and D’Andrea 2016). For details of the formulation see appendix 1-2.

We must add that until now HOIs and evolutionary dynamics have not been considered together. The role of HOIs and their links with evolutionary dynamics needs to be evaluated rigorously to understand the implications for species coexistence and our models could constitute a first step in that effort. We make our HOIs density dependent purely for mathematical simplicity, and although density-mediated HOIs could be prevalent in nature, we have no reason to presume that it is the norm. The importance of HOIs in mediating plant species coexistence has for long been suspected. There is compelling evidence for the role of soil microorganisms in stabilizing plant-plant interactions and promoting species coexistence through pervasive plant soil feedbacks (Crawford et al. n.d.). Here coexistence for pairs of plant species could be mediated by interactions with mycorrhizal species, or native microbes that play functional roles as pathogens or symbionts. For example, strong resource competition between a species pair may result in exclusion of the weaker competitor, but a third species may modulate the strength of the competitive interactions by modifying resource availability (Hawkes et al. 2005; Hinsinger et al. 2009) and reduce the fitness difference allowing for species coexistence.

### 2.3 Species coexistence in higher-order competition models with and without intraspecific variation

Using the three-way interactions community model (see section 2.2 above), we assess the influence of intraspecific trait variation on species coexistence. We examine analytically and compare species richness in this multispecies community model with and without intraspecific variation. For mathematical simplicity, in this section, we assume that intraspecific variation is same for all the species in the community such that 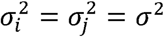. Based on strictly pairwise and three-way interactions in a diverse community, Bairey et al. (2016) derived an upper bound for species richness. Accordingly, a diverse multispecies community with pairwise as well as three-way interactions will follow (appendix 3):

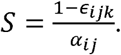

Hence the ratio of species richness with and without intraspecific variation (see appendix 3) will follow:

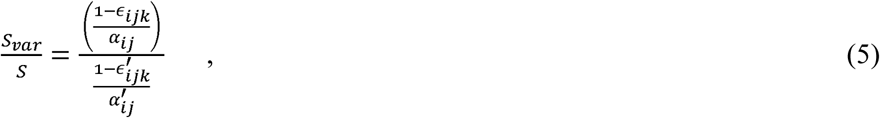

Where 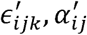 are three way and pairwise interactions without intraspecific variation, *i.e.*, 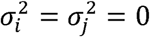 and *S*_*var*_ are s are species richness in the community with and without intraspecific variation respectively. Bairey et al. 2016 derived the expression of species richness *S* based on competition kernels with independent random variables. Their HOI terms were randomly drawn from a uniform distribution. Our equation 5 is based on gaussian competition kernels that are determined by average trait values of species. The trait values of species that are picked in starting a community are however drawn randomly from a uniform distribution (see section 2.4). The strength of competition between species is then dependent on the starting trait values. Hence, our derivation of equation 5 still holds and is a subset of the larger case presented in Bairey et. al 2016. We analyse the results from simulations of our model with this derived analytical solution of species richness, with and without intraspecific variation (see results).

### 2.4 Simulations of the community model with higher-order interactions

We assessed the effect of different levels of intraspecific trait variation on community structure and species coexistence using data generated from simulations of our community model. We simulated both trait dynamics and population dynamics resulting from equations (3) and (4). Initial species number for the start of each simulation was 40. All the 40 species were randomly assigned an initial trait value within −0.5 to 0.5 in the trait axis. Outside this trait regime, fitness value of a species will be extreme and growth rate will be negative. Effectively, this strict criterion qualitatively means that outside this trait boundary resource acquisition by a species is too low to survive and have positive growth rate. We carried out 45 replicate simulations for each level of intraspecific variation. We also simultaneously tested the influence of the width of the competition kernel, which signifies the strength of pairwise interaction, using a full factorial design where all possible combinations of intraspecific variation and interaction strength were examined for their influence on species coexistence. In all our simulations, heritability 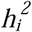 of the trait for all species was fixed at 0.1.

We evolved our community for a maximum of 1×10^4^ time points, but we concluded each simulation when the community had reached a stable state. We assumed that the community attained a stable state if the ratio of minimum value of the entropy of the community given by, – Σ*N*_*i*_ log (*N*_*i*_), at two different time points, 500 units apart (Δ*t* =500), remained bounded within 10^-5^. This condition was checked when the community had evolved for more than 5×10^3^ time points. If this condition was not met, we kept the simulation going for another 5×10^3^ time points before checking for the same condition. This condition was however met at almost every simulation indicating the tendency for convergence toward stable species density values.

#### 2.4.1 Levels of width of the competition kernel and intraspecific variation

The width of the competition kernel *w*, (see appendix 2) was varied from 0.2 through 0.45 with increments of 0.05. For each *w*, three different levels of intraspecific variation were tested in a fully factorial manner (6 different *w* values × 3 different *σ*^2^ values × 45 replicates). Specifically, for each *w*, intraspecific variation for each of the 40 species in the community was randomly sampled from a uniform distribution with three different levels: a) low variation: *σ*^2^ = [0.0006,0.003]; b) intermediate variation: *σ*^2^ = [0.003,0.009]; and c) high variation: *σ*^2^ =[0.01,0.05]; (See Table 1, for parameters used).

**Table 1:**
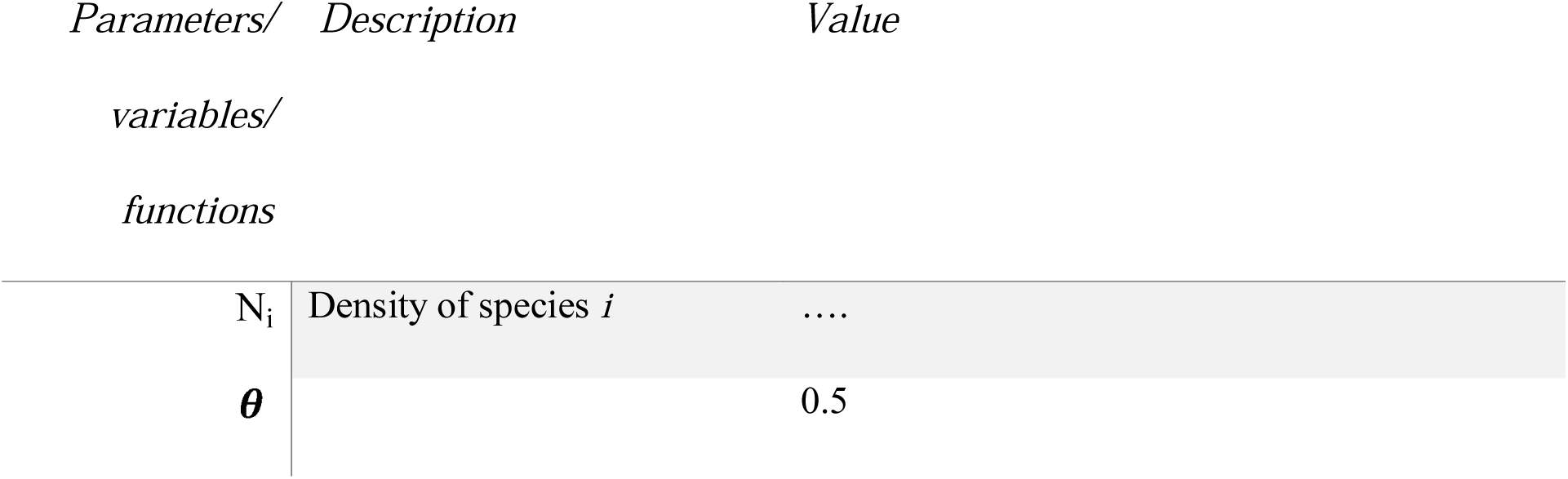
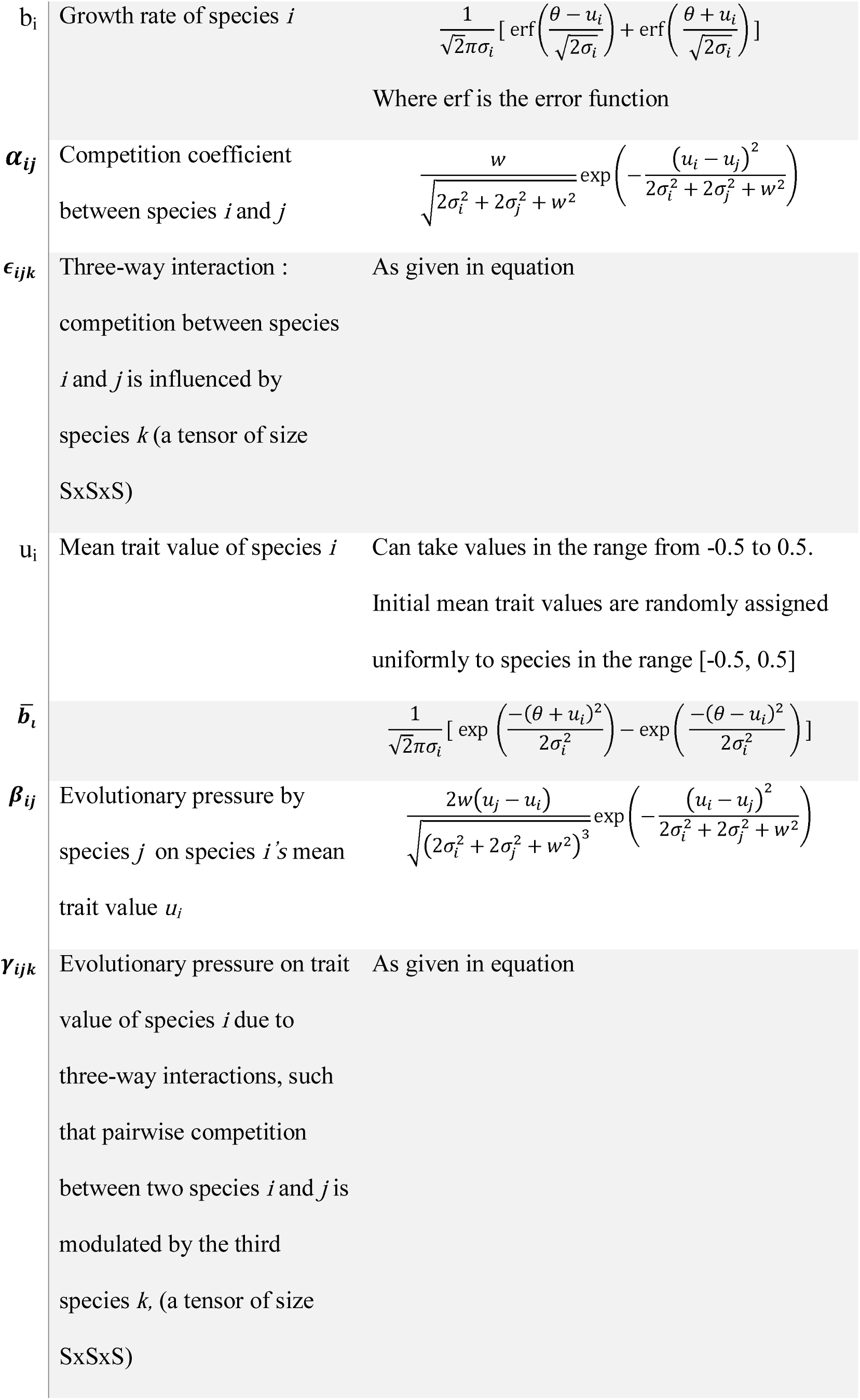
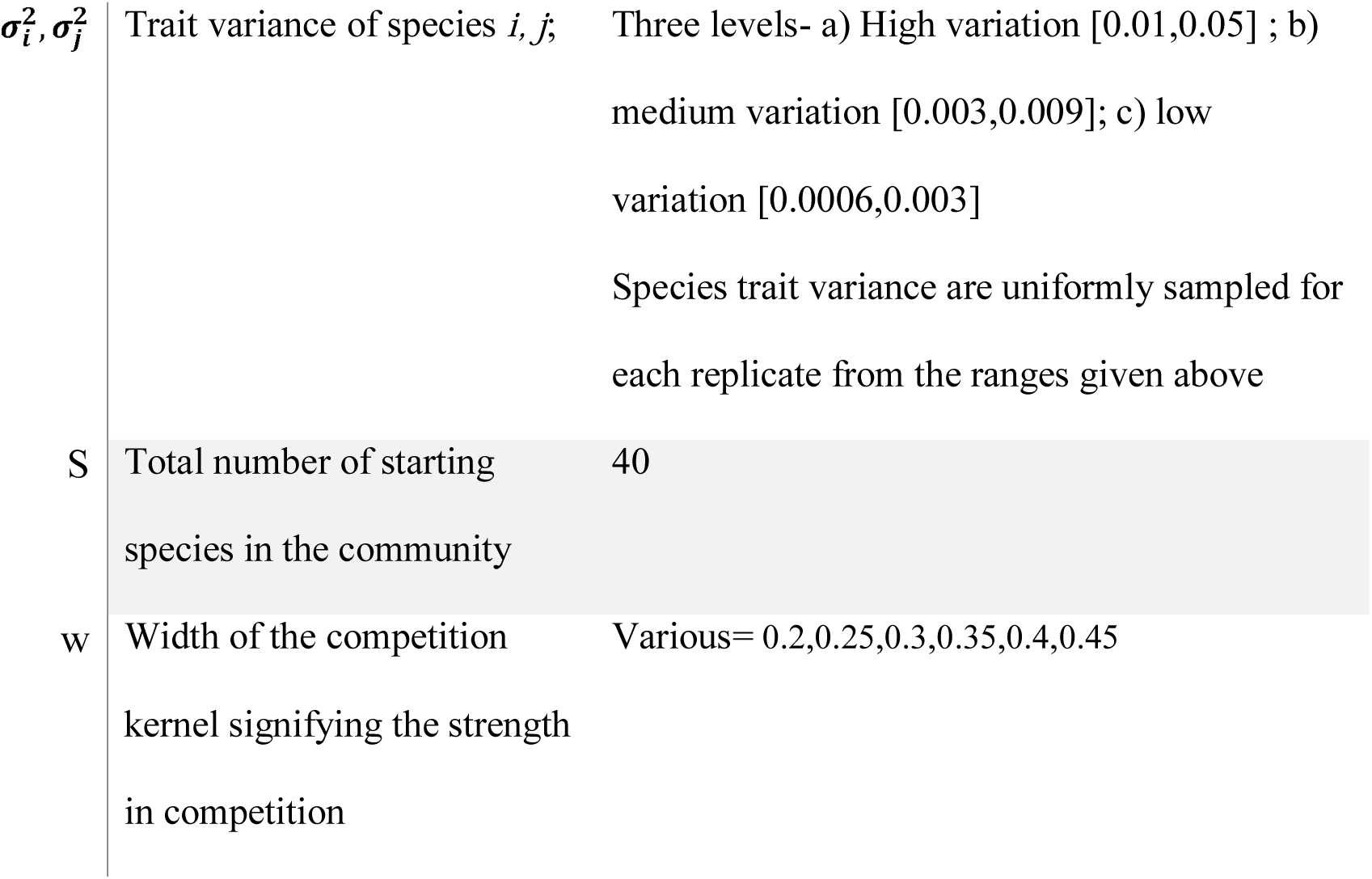
List of parameters, variables and functions of the model and their respective description and values.

### 2.5 Trait clustering

Theoretical models have suggested that species coexisting together tend to spread more evenly along a trait axis than expected (Barabas and D’Andrea 2016; D’Andrea and Ostling 2016*a*). However, empirical studies have shown that it is possible for species clusters to emerge along a trait axis (Segura et al. 2011; Vergnon et al. 2012). Here, we use a quantitative metric to evaluate the effect of intraspecific variation on the patterning of traits in the trait-axis. We measured trait similarity among coexisting species by measuring the coefficient of variation (CV) of adjacent trait means (D’Andrea and Ostling 2016*a*). High values of CV would indicate clustering of trait means of species in the trait axis while lower CV values would indicate even spacing of traits. In addition, we also compared results from trait clustering in the presence and absence of HOIs (see appendix 4).

### 2.6 Stability and robustness measures of species coexistence

Stability of our community model with higher-order interactions was measured by calculating the Jacobian at equilibrium. Specifically, the Jacobian of our dynamical system at a given point is (see appendix 5):

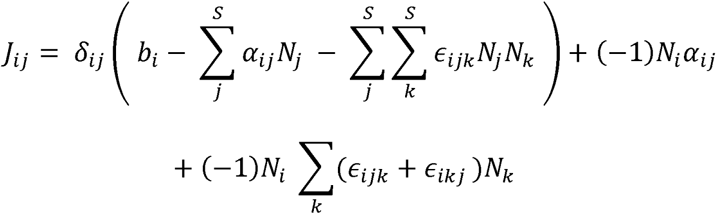

where, *δ*_*ij*_ is the Kronecker delta. At the end of our simulations, it is possible that all the species coexist, but for the community to be locally stable, the eigenvalues of the Jacobian at that point must all be negative. Thereafter, we measured the average robustness of the community by taking the geometric mean of the absolute values of the eigenvalues of the Jacobian (May 1973) (see appendix 3). Alternatively, one could calculate average robustness by the determinant of the Jacobian and that would also yield the same quantity as taking the geometric mean of the absolute values of the eigenvalues of the Jacobian. Specifically, this quantity measures the average return times in response to environmental perturbation for each of the species in the community. For each replicate simulation of each level of intraspecific variation, we calculated the average community robustness as the measure to evaluate how intraspecific variation affected robustness of species coexistence. Here, low values of average community robustness indicate lower stability.

## 3. Results

### 3.1 Analytical solution for the three-way competition model with and without intraspecific variation

We found that communities with higher intraspecific variation resulted in greater numbers of coexisting species than communities that had no intraspecific variation (Fig. 3). At low levels of intraspecific variation, the ratio of species richness with and without intraspecific variation was around 1. But as intraspecific variation increased, the ratio 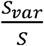 also increased significantly (Fig. 3).

**Figure 2.**
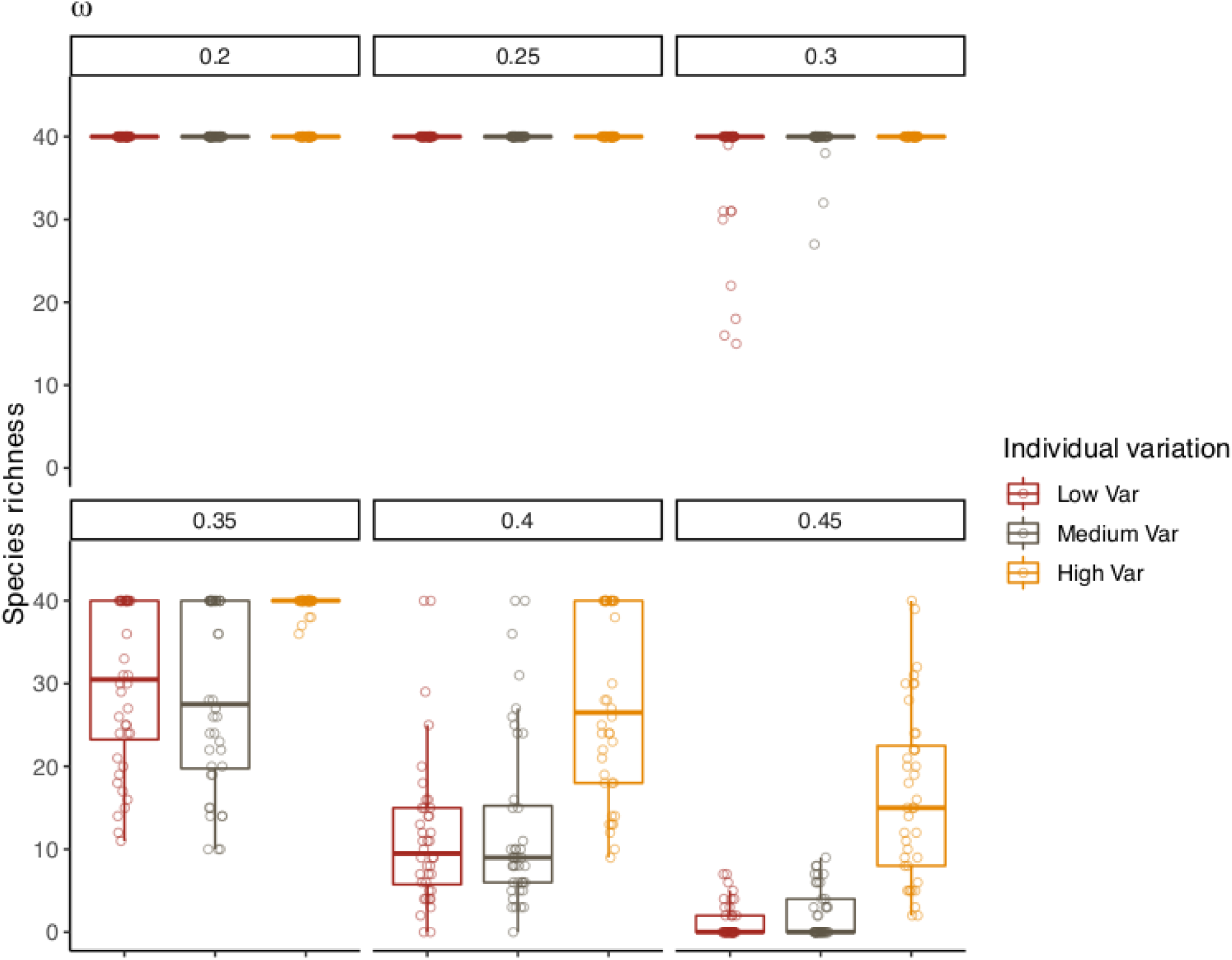
Effect of intraspecific trait variation on species richness. Boxplots denote the total number of species that coexisted at the end of the simulations for different levels of competition denoted by *w* levels. Grey colour indicates low individual variation; red colours represent medium intraspecific variation; brown colour denotes high level of intraspecific variation.

**Figure 3.**
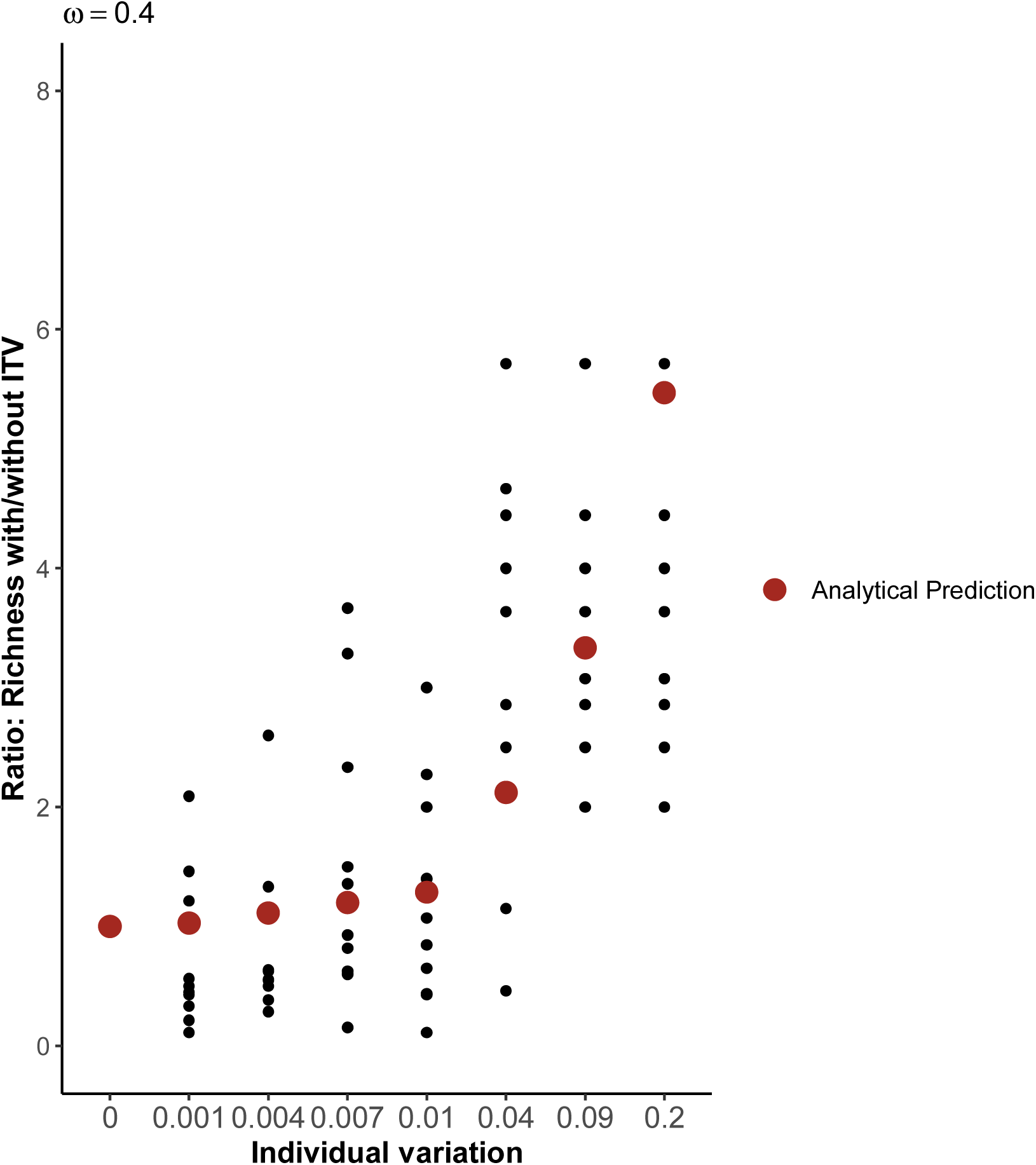
Effect of intraspecific trait variation on ratio of species richness with and without intraspecific variation. Red color dots denotes the analytical prediction of the model as given by equation (5). Black dots represent replicate model simulations for each level of intraspecific variance. Note that there is a shift in the ratio of species richness with to without intraspecific variation (ITV) once intraspecific trait variation increases above 0.01. This denotes that number of species coexisting increases significantly after intraspecific variation increases above the threshold of 0.01. This simulation and analytical prediction were shown for *w* = 0.4.

### 3.2 Effect of intraspecific variation and strength in competition on species coexistence

We found that (see above) with increases in intraspecific variation, the numbers of coexisting species increased. When we tested the interaction between competition and intraspecific variation, we found that at low levels of competition *w*, the effect of intraspecific variation on species coexistence was minimal, particularly for *w* = 0.2 and *w* = 0.25. But as the intensity of competition increased, we observed intraspecific variation had a stabilizing effect on species coexistence. At high levels of competition *w*, high intraspecific variation allowed a greater number of species to coexist on the trait axis (Fig. 2, Fig. 3).

### 3.3 Trait clustering

The coefficient of variation (CV) of trait values increased as intraspecific variation increased in the presence of HOIs, only for certain values of strength of competition (Fig. 4). Particularly, in comparison to low intraspecific variation, high intraspecific variation across different competition levels resulted in high CVs of trait values (Fig. 4). In the absence of HOIs, however, trait clustering decreases as intraspecific variation increased.

**Figure 4.**
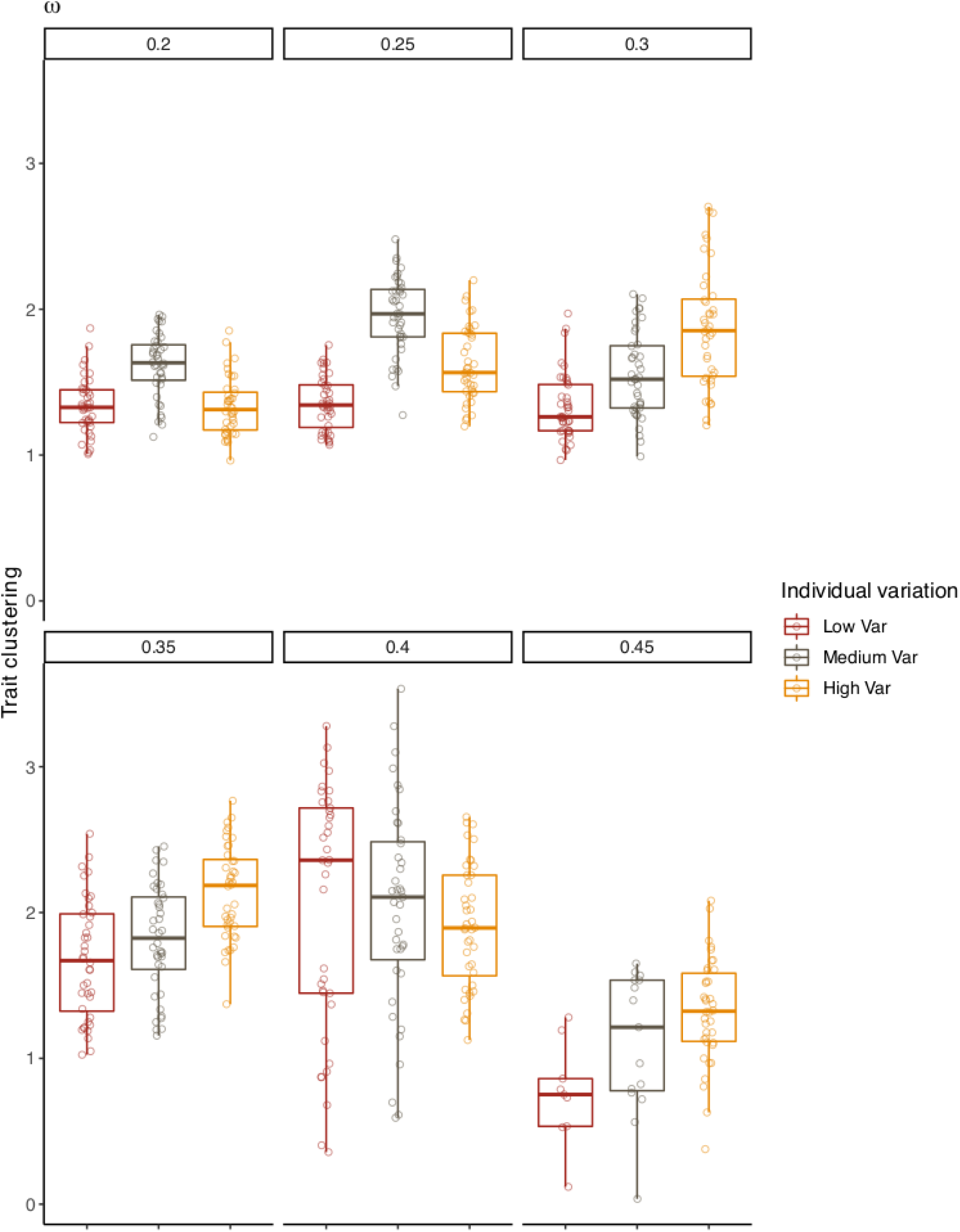
Effect of intraspecific variation on trait convergence for different levels of strength in competition, *w*. Note that trait convergence increases substantially with increasing levels of intraspecific variation across different *w*. Grey colour indicates low individual variation; red colours represent medium intraspecific variation, brown colour denotes high level of intraspecific variation.

### 3.4 Robustness of species coexistence

With increases in intraspecific variation, average robustness of the community decreased. The community became less robust to external perturbation with increasing intraspecific trait variation when compared with a community where intraspecific variation was low (Fig. 5).

**Figure 5.**
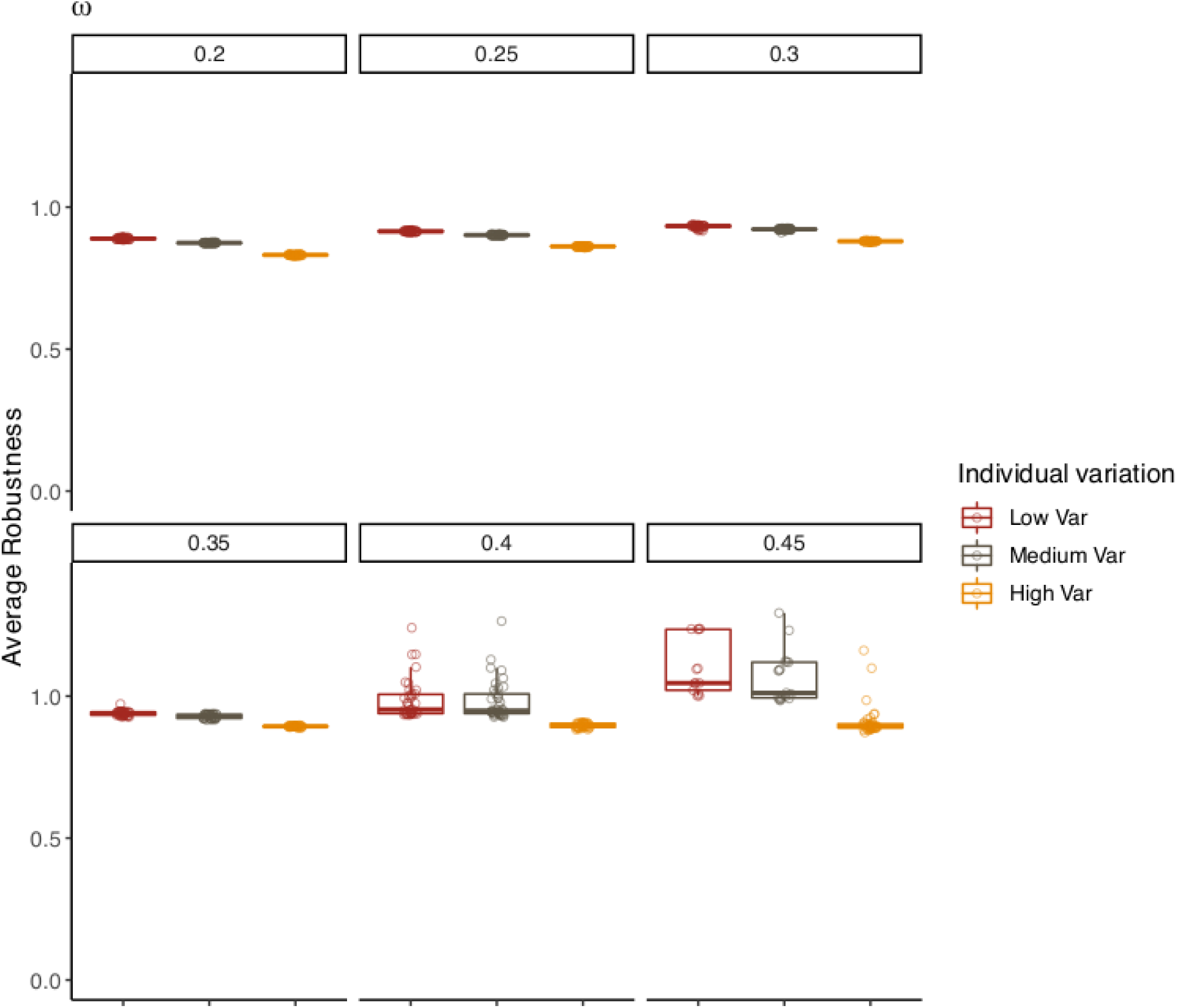
Effect of intraspecific variation on average community robustness for different levels of competition strength, *w.* Note that lower value of average robustness signifies greater community stability. With increases in intraspecific variation, average community robustness decreased across different levels of strength in competition, *w*.

## 4. Discussion

The importance and consequences of high intraspecific variation for species diversity and community structure is intensely debated (Clark 2010*b*; Clark et al. 2010; Violle et al. 2012), with contrasting findings being reported. Some studies have shown that the ecological and evolutionary consequences of individual variation are to weaken species coexistence (Barabas and D’Andrea 2016; Hart et al. 2016). Others have argued that high levels of intraspecific variation indicate that species differ primarily in the way individuals within species respond to environmental variation along multiple hidden niche dimensions. Such variable individual responses ensure that intraspecific effects are stronger than interspecific effects, a condition needed for stable species coexistence (Chesson 2000; Barabás et al. 2016). However, the nature of competitive interactions appears to be critical in determining the consequences of intraspecific variation. We investigated the ecological and evolutionary effects of intraspecific variation on coexistence in communities with both pairwise and HOIs and found strong evidence for the stabilizing effect of intraspecific variation on species coexistence.

The assumption that pairwise interactions between species are sufficient to describe competition in a community is ubiquitous in coexistence theory (Levine et al. 2017). Here, strong competition (e.g., for shared limiting resources) between pairs of species would drive species apart in niche space, structure communities, and maintain diversity. However, there is little evidence that the observed species-level differences in mean demographic rates (Condit et al. 2006) or resource use are sufficient to explain species coexistence (John et al. 2007). Here, we found strong stabilizing effect of individual variation in structuring patterns of species coexistence, provided that species interactions are mediated through HOIs.

In mechanistic models of competition where the underlying biology is modelled explicitly, HOIs can emerge subsequently in the process (Abrams 1983; Letten and Stouffer 2019). Where HOIs have been explicitly modelled in phenomenological ecological models, they act as a stabilizing factor in maintaining species diversity (Bairey et al. 2016; Grilli et al. 2017). We modelled the evolution of a trait that dictates competitive ability between species and introduced higher order competitive interactions where pairwise interactions were modulated by the density of a third species. Consistent with earlier studies on the role of HOIs (Wilson 1992; Bairey et al. 2016; Grilli et al. 2017) we found that such interactions greatly stabilize the dynamics of species in the community. Expectedly, purely pairwise interactions led to lower numbers of coexisting species as the strength of pairwise competitive interactions increased (Bairey et al. 2016) (Fig A3). In addition, as intraspecific variation increased, analogous to the results in Barabas and D’Andrea (2016), species richness in pairwise community decreased significantly (Fig A3).

A strong competitor in the trait axis can negatively affect the growth of inferior competitors. This results in a disproportionately higher abundance for the dominant competitor compared to inferior one. However, our results suggest that with the introduction of three-way interactions, this dominance of the competitively superior species is significantly reduced due to the presence of the third species, leading to proportionately similar densities for all the three species (Fig.1, Fig. A1). Our eco-evolutionary model that includes HOIs leads to stable coexistence of almost all distinct phenotypes, particularly when competitive interactions are weak. With increases in the strength of pairwise competition, higher heritable individual variation in the phenotypes stabilized ecological dynamics and led to higher numbers of coexisting species. HOIs that could emerge in species-rich competitive systems have not been well explored in the context of species coexistence (Saavedra et al. 2017). Although, empirical studies on quantifying HOIs in natural ecosystems is exceedingly difficult (Mayfield and Stouffer 2017), ignoring such interactions would limit fundamental understanding of the mechanisms behind species coexistence in complex communities.

Our results show that greater levels of intraspecific variation can lead to higher species richness, but this effect was more prominent when pairwise competition was strong (w >0.25) (Fig. 2-3). Earlier studies have shown that the numbers of species that coexists stably in eco-evolutionary models incorporating purely pairwise interactions are always less than the number of species that coexists in the absence of evolutionary dynamics (Edwards et al. 2018). With sufficient intraspecific variation, a species can evolve into a uninvasible phenotype that can lead to significant increases in its density. Consequently, the species with uninvasible phenotypes could easily displace other species in the community (Barabas and D’Andrea 2016; Edwards et al. 2018). However, with the inclusion of three-way HOIs and sufficient intraspecific variation the increase in density of superior species is significantly limited, resulting in a higher number of coexisting species. With purely pairwise interactions, eco-evolutionary models with higher intraspecific trait variation would lead to greater overlap in the trait axis and species would limit other species more than they limit themselves. Consequently, the number of species coexisting with high intraspecific variation decreases substantially (Fig A3). Under pure pairwise competition, species coexistence will be disrupted as the strength of competition increases, given a particular level of intraspecific variation. Similarly, as individual variation increases, given a particular level of competition, species coexistence will again be disrupted. Selection should therefore promote trait divergence over evolutionary time scales. Since, in our simulations we begin with saturated communities, as heritable individual variation increases species will tend to evolve away faster from one another. But in doing so, they will encounter other species at other points in the trait axis more often than when individual variation is low. Consequently, coexistence between species will decrease in the case of high individual variation. Indeed, we do observe such a result, when interactions are purely dominated by pairwise interactions. As individual heritable variation increased in the pairwise interaction case (HOIs are zero), trait clustering decreased leading to a smaller number of species that eventually coexisted (Fig A3-4).

Trait patterning varies widely, often conforming to even spacing or sometimes displaying extensive overlap (Götzenberger et al. 2012; Siefert 2012; Vergnon et al. 2012; D’Andrea and Ostling 2016*b*). In our eco-evolutionary model, where competition between species includes both pairwise and HOIs, increases in trait variation led to trait clustering (Fig. 4). Lotka-Volterra models dominated by only pairwise interactions generally support the idea that species tend to distribute more evenly along a trait axis than expected by neutral evolution for the given trait (Barabás et al. 2012; Barabas and D’Andrea 2016). This is mostly due of the underlying competition kernel and the fitness function that is generally used. Because, of the Gaussian competition kernel and the rectangular fitness function, in comparison to the species at the centre of the trait axis, species at the extreme ends of the axis will have a slight fitness advantage. This is due to the fact that species at the ends of the trait axis compete only in one direction in contrast to the species at the centre of the axis where competition is bi-directional. Expectedly, in such a scenario, due to fitness maxima at the ends of the axis, species would be displaced with respect to each other in order to minimize competitive overlap. When coupled with high heritable variation, evolution would be faster compared to when there is low heritable variation, and we should observe greater trait divergence in the former case. Indeed, in purely pairwise competitive community, we found that as intraspecific variation increased, trait convergence decreased substantially (Fig A4).

A biologically realistic case that can evaluate the sensitivity of our results and simultaneously tackle the scenario of fitness peaks at the ends of the trait axis would be by modelling a fitness function that is quadratic, such that fitness decreases unimodally towards the extremes of the trait axis. Using such an approach, however, did not alter our results (Fig A6-7). In fact, regardless of the fitness function, in the presence of HOIs, trait clustering still occurs (Fig A7). When HOIs come into play, trait divergence due to strong pairwise competition is no longer necessary as fitness loss is stabilized by a third species, and species could still persist and evolve while retaining considerable overlap in the trait axis. In other words, with the addition of HOIs, the even spacing is decreased because the third species attenuates the inhibitory or the displacing effect of the dominant species in the pairwise interaction community, thereby maintaining stable coexistence even under high trait overlap (and thus overhauling the ‘limiting similarity principle’) (Bairey et al. 2016). When high-intraspecific variation is strictly heritable, this pattern of trait clustering becomes more evident as species tend to converge on the trait axis (Tobias et al. 2014). Indeed, as intraspecific variation increased, across strength of competition, trait convergence increased, although this pattern was more or less consistent with varying strength of competition (Fig A4).

Understanding community stability when eco-evolutionary dynamics are at play is generally difficult. However, recent theory suggests that an ecological equilibrium might not be stable when different aspects of evolutionary timescales are taken into account (Patel et al. 2018). Here, our model results on community robustness are based on a time point that might not be at ecological or evolutionary equilibrium. Nevertheless, our results suggest that higher intraspecific variation leads to slightly less-robust species coexistence in the presence of HOIs (Fig. 5). This means, that with higher intraspecific trait variation, communities become less robust to external environmental perturbation (Barabas and D’Andrea 2016). High intraspecific variation led to faster evolutionary dynamics and more species coexisting together in a uni-dimensional trait axis only in the presence of HOIs. Consequently, traits of persisting species had fewer locations in the trait axis that were advantageous for average community stability. Contrastingly, studies that focused on pairwise interactions alone, have reported the stabilizing effect of higher intraspecific variation on community robustness (Barabas and D’Andrea 2016).

Although our results demonstrate the power of HOIs in structuring patterns of species coexistence, they do not yet link with insights gained from modern coexistence theory (MCT) (Chesson 2000). MCT hinges on the mechanisms that stabilize or equalize fitness differences among coexisting species (Adler et al. 2007). A recent study on linking HOIs with MCT shows that species coexistence is possible when HOIs alleviate interspecific competition between species to a greater extent than the decrease in intraspecific competition (Singh and Baruah 2019). Regardless of large fitness differences where pairwise species coexistence was impossible, it was suggested that HOIs can stabilize species coexistence provided, *ϵ*_*ijk*_ < *ϵ*_*iik*_where *i*≠*j* This means, if a third species *k* intensifies intraspecific competition more than interspecific competition, species coexistence is possible even when there are large fitness differences (Singh and Baruah 2019). With the rectangular fitness function in our model, we ensure that at any time point fitness differences between any two species is 1. Hence, when niche overlap increases due to increases in intraspecific variation, and when interactions between species are predominantly pairwise, the probability of species coexistence of a species pair decreases. The only way possible for density-mediated HOIs to promote coexistence even when there is high overlap is when intraspecific competition is further strengthened by a third species. Indeed, this is what we also observe from our results (Fig A5). However, investigations on linking these dynamics to the concepts of MCT would be necessary to further confirm our exploratory results.

We are getting closer to understanding how species richness is maintained despite differences in competitive abilities. Density mediated HOIs as modelled here, intensifies competition rather than alleviating pairwise competition, and could stabilize fitness differences by increasing pairwise intraspecific effects relative to interspecific effects (Fig A5). Even when competition was further intensified species coexistence was still possible implying the stabilizing effects of HOIs, with strength of intraspecific HOI being higher than interspecific HOIs (Fig. A5). This suggests that, in terms of MCT, HOIs as modelled here strengthened intraspecific effects more than it strengthened interspecific effects (Fig. A5) (also see Singh and Baruah 2019). HOI terms could also be positive, indicating the facilitative effects of HOIs on species pairwise interactions. It has been shown recently, however, that positive HOIs can lead to infeasible invasion growth rates (Baruah and Singh, 2019). However, further developments on this aspect is needed to understand the effect of positive HOIs on species coexistence.

Another mechanism of HOI, which has not been tested either empirically or theoretically, that could emerge when there are more than two competitors is trait-mediated HOIs (Levine et al. 2017). For instance, trait-mediated HOI could emerge when a third competitor induces a plastic change in the trait of a species in direct competition with another species. Such a trait-mediated change could intensify or alleviate pairwise competitive interactions, and in turn could either destabilize or stabilize species coexistence respectively. However, the results from such an approach might differ from the results of density-mediated HOI, particularly because of how plasticity of the trait is modelled in relation to other competitors. If plasticity is modelled as adaptive to changes in competitor’s trait, then such trait change could stabilize coexistence. However, in a saturated community this could become complicated because of multiple competitors that could potentially lead to a change in the trait in a direction that might not be favourable for species coexistence.

Our work demonstrates the importance of within species variation in maintaining species coexistence. The significance of our results demonstrates that HOIs in competition and coexistence studies should not be ignored. Thus an important next step would be to characterize higher order interactions as well as individual variation in relation to capturing variation in fitness in a diverse species community.

## Acknowledgements

We thank David Vassuer and another anonymous reviewer whose comments substantially improved the manuscript.

