## appendix for "Intraspecific variation promotes species coexistence and trait clustering through higher order interactions"

### 1. Lotka-Volterra pairwise interaction model

We model the eco-evolutionary dynamics of S species that competes for resources in a community. Competition and growth of an individual of a species is dictated by a quantitative trait $z$ [5] . An individual of a species $i$ can be described by its quantitative trait value $z$ in a uni-dimensional trait axis. Accordingly, the distribution of the quantitative trait $z$ is Gaussian, $p(z)$, with mean $\mu_i$ for species $i$ and variance $\sigma_i$. If $N_i(t)$ is the number of individuals of species $i$ in time t, then $N_i(t)p_i(z,t)dz$ is the population density of species $i'$s individuals with trait value between $z$ and $z+dz$ [5].

Following Barabas et al, 2016, the per capita Lotka-Volterra growth rate for $S$ species in a competitive community dominated by pairwise interactions can be written as:

$$r(\vec{N},\vec{p},z) = b(z) - \sum_{j=1}^{S} N_j \int \alpha(z,z')p_j(z')dz', \tag{1}$$

where $b(z)$ is the instrinsic growth rate of trait $z$ and $\alpha(z,z')$ is the competition function that captures competition between trait $z$ and $z'$. The per captita

growth rate in equation (1) is directly dependent on trait $z$. Competition among individuals of different species in the community also depends solely on the trait $z$. Further, the summation term in equation (1) ensures that every species in the community has an effect on the growth of species $i$ in the community. Our model is in the quantitative genetic limit [4] in the sense that the trait distribution $p(z)$ remains normal and only the mean of the distribution $\mu_i$ is subjected to selection due to competition and growth. Thus the variance and shape of $p(z)$ remains undisturbed.

### 2. Lotka-Volterra higher order interactions

With this pairwise Lotka-Volterra quantitative model, we introduce higher order interactions in equation (1) in the following way:

$$r(\vec{N},\vec{p},z) = b(z) - \sum_{j=1}^{S} N_j \int \alpha(z,z')p_j(z',t)dz' - \sum_{k=1}^{S}\sum_{j=1}^{S} N_k N_j \int \epsilon_k(z,z')p_j(z',t)dz', \tag{2}$$

Where $\epsilon_k(z,z')$ is the pair-wise interaction term of individuals of two different traits $z$ and $z'$ (in the same way as $\alpha(z,z')$) . However,the pairwise interaction between two individuals is modulated by the density of a third species $k$. The double summation ensures that each species modulates the pairwise interaction between two other species.

$\alpha(z,z')$ is the Gaussian competition kernel in (1) such that individuals with very similar trait will compete strongly whereas individuals that are far apart in trait values $z$ will compete weakly, given as:

$$\alpha(z,z') = \exp\big(\frac{-(z-z')^2}{\omega^2}\big),$$

Further,

$$\epsilon_k(z,z') = N_k \exp\big(\frac{-(z-z')^2}{\omega^2}\big)$$

where $\epsilon_k(z,z')$ scales with the density of the species $k$[8] .

Growth rate of an individual with trait value $z$ is given by $b(z)$ which is a rectangular function along the trait axis given as:

$$b(z) = \begin{cases} 1, & \text{if } -\theta \geq z \geq \theta \\ 0, & \text{otherwise,} \end{cases}$$

Where $\theta$ is the limit of the trait axis such that any individual which has a trait value outside of the range of $[-\theta, \theta]$ will have zero growth.

With this growth equation, the dynamics of species $i$ can be written as :

$$\frac{dN_i(t)}{dt} = \int r_i(\vec{N}, z, t) p_i(z, t) dz, \tag{3}$$

Expanding equation (3) gives:

$$\frac{dN_i(t)}{dt} = N_i(t) \int \Big( b(z) - \sum_{j=1}^{S} N_j \int \alpha(z, z') p_j(z') dz' - \\ \sum_{k=1}^{S} \sum_{j=1}^{S} N_k N_j \int \epsilon_k(z, z') p(z, z') dz' \Big) p_i(z, t) dz,$$

$$\frac{dN_i(t)}{dt} = N_i(t) \Big( \int \big( b(z) p_i(z, t) dz \big) - \sum_{j=1}^{S} N_j \int \int \alpha(z, z') p_j(z', t) p_i(z, t) dz' dz - \\ \sum_{k=1}^{S} \sum_{j=1}^{S} N_k N_j \int \int \epsilon_k(z, z') p_j(z', t) p_i(z, t) dz dz' \Big),$$

Finally simple algebra of the above equations would lead to

$$\frac{dN_i(t)}{dt} = N_i(t) \Big( b_i(t) - \sum_{j=1}^{S} \alpha_{ij}(t) N_j(t) - \sum_{k=1}^{S} \sum_{j=1}^{S} \epsilon_{ijk}(t) N_j(t) N_k(t) \Big), \tag{4}$$

Where,

$$\alpha_{ij}(t) = \int \int \alpha(z, z') p_j(z', t) p_i(z, t) dz' dz = \frac{\omega}{\sqrt{2\sigma_i^2 + 2\sigma_j^2 + \omega^2}} \exp \big( \frac{-(\mu_i(t) - \mu_j(t))^2}{2\sigma_i^2 + 2\sigma_j^2 + \omega^2} \big). \tag{5}$$

The double integral is the weighted sum of interactions between individual of species $i$ which has $p_i(z)$ as its trait distribution and species $j$ which has $p_j(z')$ as its trait distribution. This term quantifies the competition species $i$ faces from species $j$. And,

$$b_i(t) = \int b(z)p_i(z,t)dz = \frac{1}{2}\Bigg[erf(\frac{\theta - \mu_i}{\sqrt{2}\sigma_i}) + erf(\frac{\theta + \mu_i}{\sqrt{2}\sigma_i})\Bigg]. \tag{6}$$

$$\epsilon_{ijk}(t) = \int\int \epsilon_k(z,z')p_j(z',t)p_i(z,t)dzdz' = N_k(t)\frac{\omega}{\sqrt{2\sigma_i^2 + 2\sigma_j^2 + \omega^2}} \exp\big(\frac{-(\mu_i(t) - \mu_j(t))^2}{2\sigma_i^2 + 2\sigma_j^2 + \omega^2}\big). \tag{7}$$

The above term captures the three-way strength of interaction. Pairwise competition between species $i$ with mean trait $\mu_i$ and species $j$ with mean trait $\mu_j$ is influenced by the density of another species $k$ [8]. Finally the dynamics of the trait of a species in response to growth as well as in response to interspecific competition due to pairwise interactions among species and higher-order interactions can be formulated as ,

$$\frac{d\mu_i(t)}{dt} = h_i^2\sigma_i^2\frac{\partial}{\partial\mu_i}(\frac{1}{N_i}\frac{dN_i(t)}{dt}), \tag{8}$$

where, $h_i^2$ is the fraction of heritable variation of the phenotype $\mu_i$ of species $i$. In our simulations we fixed the heritable variation to 0.1 for all the species. This can thus be further expanded to :

$$\frac{d\mu_i(t)}{dt} = h_i^2\sigma_i^2\frac{\partial}{\partial\mu_i}\Big(b_i(t) - \sum_{j=1}^{S}\alpha_{ij}(t)N_j(t) - \sum_{k=1}^{S}\sum_{j=1}^{S}\epsilon_{ijk}(t)N_j(t)N_k(t)\Big),$$

Differentiating the first term of the above equation with respect to $\mu_i$ we get:

$$\frac{\partial}{\partial\mu_i}b_i(t) = \frac{\partial}{\partial\mu_i}\Big(\frac{1}{2}\Bigg[erf(\frac{\theta - \mu_i}{\sqrt{2}\sigma_i}) + erf(\frac{\theta + \mu_i}{\sqrt{2}\sigma_i})\Bigg]\Big) = \frac{1}{\sqrt{2\pi}\sigma_i}\Bigg[\exp\big(\frac{-(\theta + \mu_i)^2}{2\sigma_i^2}\big) - \exp\big(\frac{-(\theta - \mu_i)^2}{2\sigma_i^2}\big)\Bigg],$$

where $erf$ is the error function. Thus:

$$\bar{b_i}(t) = \frac{1}{\sqrt{2\pi}\sigma_i}\Bigg[\exp\big(\frac{-(\theta+\mu_i)^2}{2\sigma_i^2}\big) - \exp\big(\frac{-(\theta-\mu_i)^2}{2\sigma_i^2}\big)\Bigg].$$

Similarly,

$$\frac{\partial}{\partial \mu_i}\alpha_{ij}(t) = \frac{2\omega(\mu_j(t)-\mu_i(t))}{\sqrt[3]{2\sigma_i^2+2\sigma_j^2+\omega^2}}\exp\big(-\frac{(\mu_i(t)-\mu_j(t))^2}{2\sigma_i^2+2\sigma_j^2+\omega^2}\big) = \beta_{ij}(t)$$

.

And,

$$\frac{\partial}{\partial \mu_i}\epsilon_{ijk}(t) = N_k(t)\frac{2\omega(\mu_j(t)-\mu_i(t))}{\sqrt[3]{2\sigma_i^2+2\sigma_j^2+\omega^2}}\exp\big(-\frac{(\mu_i(t)-\mu_j(t))^2}{2\sigma_i^2+2\sigma_j^2+\omega^2}\big) = \gamma_{ijk}(t).$$

Thus

$$\frac{d\mu_i(t)}{dt} = h_i^2\sigma_i^2\Big(\bar{b_i}(t) - \sum_{j=1}^{S}\beta_{ij}(t)N_j(t) - \sum_{k=1}^{S}\sum_{j=1}^{S}\gamma_{ijk}(t)N_j(t)N_k(t)\Big). \quad (9)$$

From equation 9, the first term denotes evolutionary pressure due to growth of the trait in the trait axis, the second term $\beta_{ij}$ denotes the selection acting on the mean trait value of species $i$ due to pairwise competition with other species in the trait axis, and the third term $\gamma_{ijk}$ indicates the selection that acts on the trait due to higher-order interactions in the trait axis.

### 3. Analytical calculation of species richness with and without intraspecific variation

Bairey et al (2015) extended May's result of diversity and stability of communities with pairwise interactions to communities with higher-order interactions. They suggested that diversity of a large community can scale differently to interactions of different orders as:

$$\alpha + \frac{\epsilon}{S} + \frac{\gamma'}{S^2} + ... \leq \frac{1}{S} \quad (10)$$

where $\alpha$, $\epsilon$ and $\gamma'$ are pairwise, three-way and fourth-order interactions and $S$ is the number of species in the community. Extending this result they found that these scaling relationship can be used to estimate the optimal community size as :

$$S_{min,max} = \frac{1-\epsilon \pm \sqrt{(1-\epsilon)^2 - 4\alpha\gamma'}}{2\alpha} \tag{11}$$

We can use the result from equation (10) and (11) to derive the optimal species richness (or community size) with and without intraspecific variation. In our model fourth-order interaction is zero such that $\gamma' = 0$. Hence we can re-write the above relationship as follows:

$$S_{min,max} = \frac{1-\epsilon \pm (1-\epsilon)}{2\alpha} \tag{12}$$

which gives us the maximum and minimum number of species that could coexist given the community is dictated by both pairwise interactions and three-way higher order interactions. Thus the maximum number of species that could coexist in a large community in our model would be :

$$N_{max} = \frac{1-\epsilon}{\alpha}$$

For easier analytical calculation, we consider two simple cases : 1) a community with $S'$ species with no intraspecific variation and 2) and a community of $S$ species with intraspecific trait variation.

Hence the ratio of species richness with and without intraspecific variation will follow :

$$\frac{S}{S'} = \frac{\frac{1-\epsilon}{\alpha}}{\frac{1-\epsilon'}{\alpha'}} \tag{13}$$

where $\epsilon$ is the three-way higher order interaction for the community with intraspecific variation given as from (7) as:

$$\epsilon = N\frac{\omega}{\sqrt{4\sigma^2+\omega^2}}\exp(\frac{-d^2}{4\sigma^2+\omega^2})$$

where $d^2$ is the average trait difference between species in the trait axis. And $\epsilon'$ is the three way interaction term for the community without any intraspecific variation given as from (7) (by setting $\sigma_i^2 = 0$):

$$\epsilon' = N \exp(\frac{-d^2}{\omega^2})$$

Similarly the pairwise interaction for the community with intraspecific variation from (5),

$$\alpha = \frac{\omega}{\sqrt{4\sigma^2 + \omega^2}} \exp\big(\frac{-d^2}{4\sigma^2 + \omega^2}\big)$$

And for the community without intraspecific variation would be :

$$\alpha' = \exp\big(\frac{-d^2}{\omega^2}\big)$$

Which leads, after rearranging equation (13), to :

$$\frac{S}{S'} = \frac{\frac{1 - N\frac{\omega}{\sqrt{4\sigma^2+\omega^2}}\exp(\frac{-d^2}{4\sigma^2+\omega^2})}{\frac{\omega}{\sqrt{4\sigma^2+\omega^2}}\exp\left(\frac{-d^2}{4\sigma^2+\omega^2}\right)}}{\frac{1 - N\exp(\frac{-d^2}{\omega^2})}{\exp\left(\frac{-d^2}{\omega^2}\right)}} \tag{14}$$

And plotting $\frac{S}{S'}$ for various levels of $\sigma$ or in other words for various levels of intraspecific variation from low to high leads to Fig. 4 in main text and fig A2.

### 4. Intraspecific variation and strength in higher order interactions

From Bairey et al (2015) [1], the optimal community size is a function of the strength of pairwise interaction as well as a function of strength of higher-order interaction, given by equation (11). However, when a community exhibits higher-order interactions, does higher individual variation lead to lowering of the strength of such interactions?

In our model, the three-way interaction is given by :

$$\epsilon_{ijk} = N_k(t)\frac{\omega}{\sqrt{2\sigma_i^2 + 2\sigma_j^2 + \omega^2}} \exp\big(\frac{-(\mu_i(t) - \mu_j(t))^2}{2\sigma_i^2 + 2\sigma_j^2 + \omega^2}\big).$$

Assuming, that species $i$ and $j$ have equal trait variance such that, $\sigma_i^2 = \sigma_j^2 = \sigma^2$, we can rewrite this as :

$$\epsilon_{ijk} = N_k(t)\frac{\omega}{\sqrt{4\sigma^2+\omega^2}}\exp\big(\frac{-(\mu_i(t)-\mu_j(t))^2}{4\sigma^2+\omega^2}\big)$$

Plotting various levels of trait variation $\sigma^2$ with the strength in three way interaction reveals that strength in the three way interaction $\epsilon_{ijk}$ decreases as intraspecific variation increases (see Fig. A3).

### 5. Jacobian, stability and robustness of species coexistence

The Jacobian of a dynamical system at a given point is given as:

$$J_{ij} = \frac{\partial\big(\frac{dN_i(t)}{dt}\big)}{\partial N_j}$$

.

At equilibrium, when changes in the abundances of species is zero which is the equilibrium condition of S species coexisting, then we can write from equation (3) that :

$$\frac{dN_i(t)}{dt} = \Big(b_i - \sum_{j=1}^{S}\alpha_{ij}N_j(t) - \sum_{k=1}^{S}\sum_{j=1}^{S}\epsilon_{ijk}N_j(t)N_k\Big) = 0, \tag{15}$$

which means at equilibrium, species $i$ will follow the equation below :

$$\Big(b_i = \sum_{j}^{S}\alpha_{ij}N_j - \sum_{k}^{S}\sum_{j}^{S}\epsilon_{ijk}N_jN_k\Big). \tag{16}$$

Evaluating the Jacobian at this equilibrium point from equation (16) leads to:

$$J_{ij} = \frac{\partial\big(\frac{dN_i(t)}{dt}\big)}{\partial N_j} = \delta_{ij}\Big(b_i - \sum_{j}^{S}\alpha_{ij}N_j - \sum_{k}^{S}\sum_{j}^{S}\epsilon_{ijk}N_jN_k\Big) - N_i\alpha_{ij} - \delta_{ij}N_i - N_i\Big(\sum_{k}^{S}\epsilon_{ijk} + \sum_{k}^{S}\epsilon_{ikj}\Big) \tag{17}$$

where $\delta_{ij}$ is the Kronecker delta where

$$\delta_{ij} = \begin{cases} 1, & \text{if } i = j \\ 0, & \text{otherwise.} \end{cases}$$

Now at equilibrium,

$$\Big(b_i = \sum_{j}^{S} \alpha_{ij} N_j - \sum_{k}^{S} \sum_{j}^{S} \epsilon_{ijk} N_j N_k\Big),$$

Hence equation (17) can be re-written at equilibrium as:

$$\frac{\partial\big(\frac{dN_i(t)}{dt}\big)}{\partial N_j} = -N_i\alpha_{ij} - \delta_{ij}\Big(b_i - \sum_{j}^{S} \alpha_{ij} N_j - \sum_{k}^{S} \sum_{j}^{S} \epsilon_{ijk} N_j N_k\Big) - N_i\Big(\sum_{k}^{S} \epsilon_{ijk} + \sum_{k}^{S} 2\epsilon_{ikj}\Big). \quad (18)$$

Equation (18) is a modified community matrix that incorporates higher-order 3 way interaction [7] . From this equation at particular equilibrium time point, one can estimate eigenvalues from this Jacobian matrix and in turn can estimate stability and average robustenss of species coexistence at equilibrium.

If the eigenvalues of the equation (18) yields all negative eigenvalues, then the community at that point is stable. In other words, suppose, S species coexist at the end of a simulation run, and if the eigenvalues of the Jacobian at that time point has all S negative eigenvalues then the equilibrium is locally stable.

Thus local stability of an ecological community will be guaranteed if the Jacobian matrix in equation (19) has eigenvalues that have negative real parts. This means that any perturbation at that point will decay along its eigendirection with a rate equal to the eigenvalue. The robustness of this local stability of an ecological community is given by the determinant of the Jacobian matrix [2, 6, 3]. As explicitly formulated in Barabas et al 2015, it is given by geometric mean of the absolute values of the eigenvalues as :

$$\sqrt[S]{|\lambda_1||\lambda_2|...|\lambda_S|} = \Big(\prod_{i=1}^{S} |\lambda_i|\Big)^{\frac{1}{S}} = \exp(\frac{1}{S}\log(\prod_{i=1}^{S} |\lambda_i|) = \exp(\overline{\log}(|\lambda|))$$

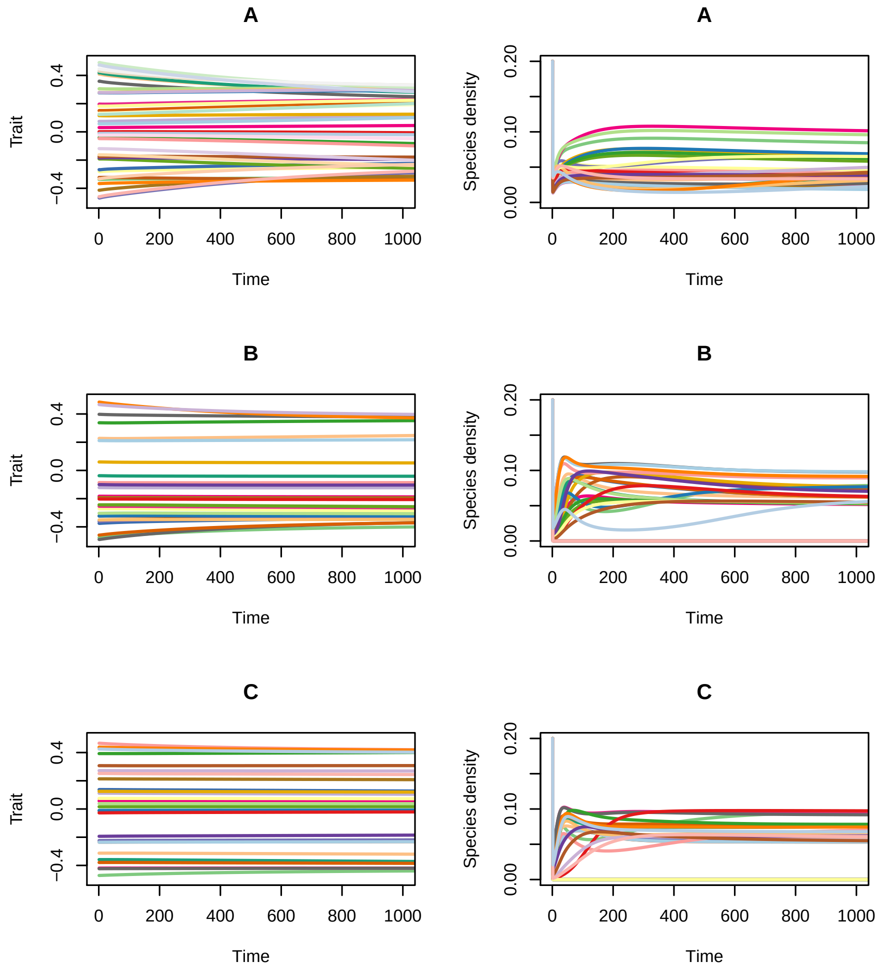


Fig.A 1: Example dynamics of trait (left column) and species density (right column) over time for three levels of intraspecific variation for a particular w = 0.3. (A) High intraspecific variation; (B) Medium intraspecific variation; (c) low intraspecific variation. Different colors represent different species.

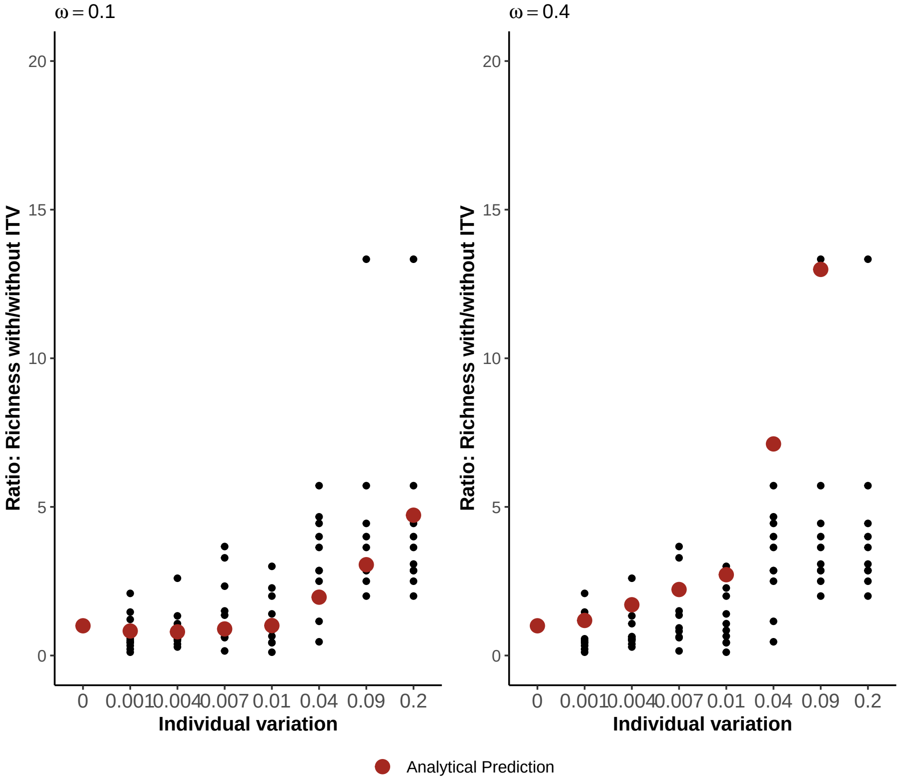


Fig.A 2: Community richness ratio with to without intraspecific variation on Y-axis and different levels of intraspecific variance on X-axis. The black dots represent replicate simulations of the model for different levels of intraspecific variation and the red dots is the analytic prediction from equation 14, for two levels of interspecific strength in competition are $w = 0.1 and 0.4$ and $(\mu_i - \mu_j) = 0.1$, $N_k = 1$

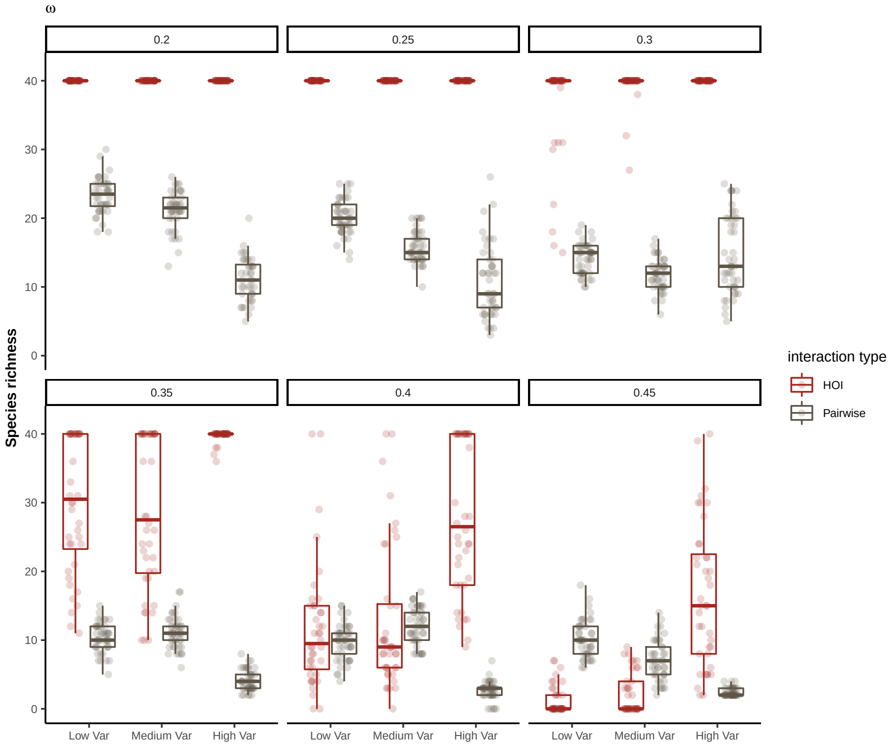


Fig.A 3: Effect of intraspecific trait variation on species richness across different levels of competition for two different interaction types : HOI and pairwise. Boxplots denote the total number of species that coexisted at the end of the simulations for different levels of competition denoted by w levels.

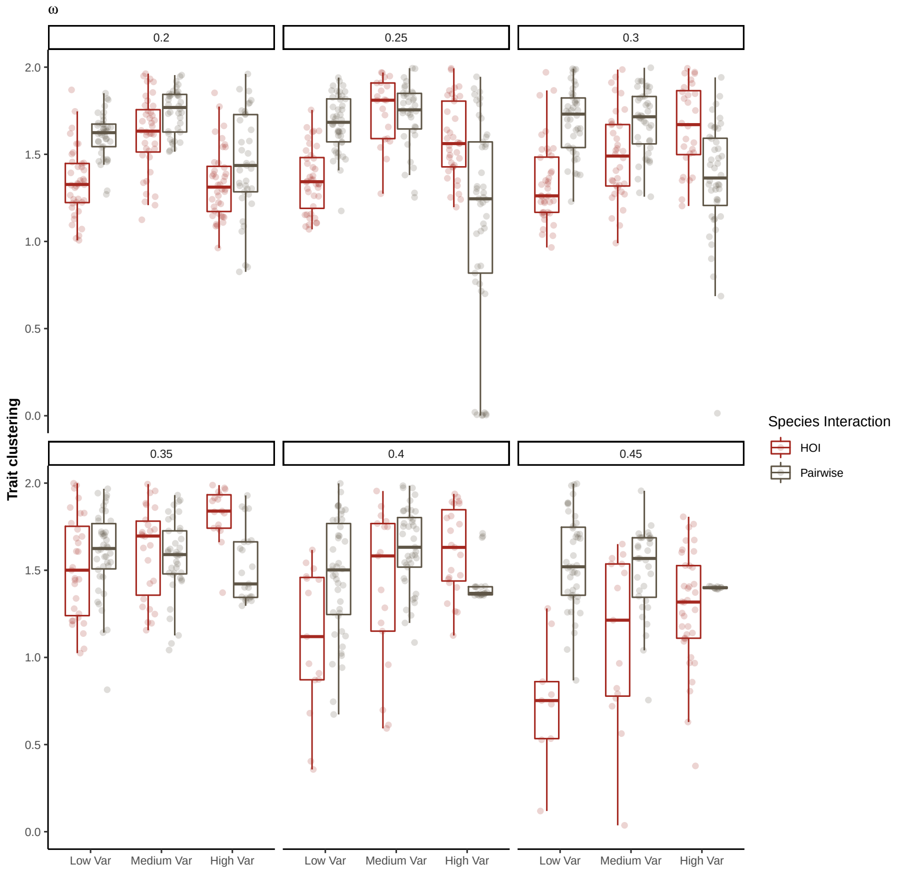


Fig.A 4: Effect of intraspecific variation on trait convergence for different levels of strength in competition, w and for two different interaction types : HOI and pairwise. Note that trait convergence increases with increasing levels of intraspecific variation for HOI interactions, whereas the pattern is opposite for pure pairwise interaction.
